# Metacognitive Reactivity and Temporal Perception: The Influence of Confidence Ratings on the Temporal Binding Window

**DOI:** 10.1101/2025.02.28.640845

**Authors:** Carolin Sachgau, Aysha Basharat, Michael Barnett-Cowan

## Abstract

The influence of confidence in assessing the perceived timing of sensory events on perceptual decision-making remains poorly understood. Metacognition—the ability to reflect on one’s thought processes — is often assessed through confidence ratings in perceptual tasks. However, accessing metacognitive information can influence task performance, a phenomenon known as the reactivity effect. This study examines metacognitive reactivity by examining the impact of confidence ratings on the temporal binding window (TBW) in two multisensory time perception tasks that use identical stimuli but require different judgments: Temporal Order Judgment (TOJ) and Simultaneity Judgment (SJ). Thirty-five participants (aged 18–39) completed TOJ and SJ tasks with and without confidence ratings via Prolific, an online participant recruitment platform. Psychometric functions were fitted to response and confidence data to determine TBW and the point of subjective simultaneity (PSS). Both tasks exhibited reactivity effects, with the effect being stronger in the SJ task. TBW significantly differed between TOJ and SJ tasks, regardless of confidence ratings. Additionally, TBWs from TOJ and SJ tasks were strongly correlated, as were the width parameters of confidence curves, suggesting shared underlying mechanisms in both perceptual and metacognitive processes. These findings highlight the utility of confidence ratings in evaluating TOJ and SJ performance, revealing both task differences and commonalities. We also discuss the potential for training interventions to enhance temporal perception, particularly in older adults. Finally, we reflect on the advantages and challenges of online data collection via Prolific, including its diverse participant pool and timing precision limitations.

## INTRODUCTION

Metacognition—the ability to reflect on one’s own thought processes—is a fundamental aspect of human cognition, including the perception of sensory event timing. It is commonly assessed through self-report measures, where participants provide subjective evaluations of their performance during or after a task. Confidence ratings are widely used to quantify metacognition (Fleming, 2024; Grimaldi et al., 2015; Yeung & Summerfield, 2012), typically on categorical (such as high vs low confidence) or continuous scales ranging from 0–100% or 50–100% (Grimaldi et al., 2015). However, the act of assessing metacognitive information can itself alter task performance, a phenomenon known as the reactivity effect (Kazdin, 1974). Several factors influence the magnitude and direction of this effect, including task difficulty, trait confidence of the participant, and even the phrasing of confidence-related questions (see Double & Birney, 2019 for a review).

Despite extensive research on metacognition in various domains, relatively few studies have examined its role in time perception (Lamotte & Droit-Volet, 2017). However, evidence suggests that metacognition influences temporal judgments. For example, increased awareness of attention’s role in time perception reduces temporal underestimation (Lamotte et al., 2012) Allan (1975) demonstrated that psychometric functions for high-versus low-confidence responses were nonparallel, indicating distinct cognitive processes based on confidence. Furthermore, Keane et al (2015) found that in a temporal order judgment (TOJ) task, confidence was positively correlated with timing precision such that when people are highly confident, their audiovisual timing judgements are more precise, a result mirrored in duration judgment tasks (Lamotte and Droit-Volet, 2017). Similarly, Akdogan and Balci (2017) showed that confidence ratings reflect awareness of both the direction and magnitude of errors in a time reproduction task. However, the impact of confidence ratings on time perception performance, particularly in terms of reactivity, remains largely unexplored. Some evidence suggests that participants can recognize and correct temporal biases when incentivized (Brocas et al., 2018), implying that metacognitive awareness could influence performance.

While time perception is often studied in a single sensory modality, the brain must integrate information across multiple senses to form coherent percepts, a process known as multisensory integration (Deroy et al., 2016). This process involves binding stimuli that belong together while segregating those that do not, despite differences in neural processing speeds. The interval within which stimuli must be presented to be perceived as simultaneous is termed the temporal binding window (TBW) (Meredith et al., 1987; Stein et al., 1988). The TBW is typically measured using TOJ tasks, where participants judge which of two stimuli appeared first, and simultaneity judgment (SJ) tasks, where participants determine whether stimuli occurred simultaneously (van Eijk et al., 2008). Despite the importance of multisensory integration, the role of confidence in multisensory perceptual tasks has been understudied (see Maynes et al., 2023; Mims et al., 2020) and the reactivity effect in such tasks remains largely unexplored. Emerging evidence suggests that confidence judgments vary across sensory modalities. For instance, in a temporal bisection task, participants exhibited greater metacognitive awareness in audiovisual compared to unisensory conditions (Cropper et al., 2024). Given the inherently multisensory and dynamic nature of real-world environments, it is critical to study metacognition in multisensory contexts to fully understand its role in time perception. To our knowledge, no study has directly investigated the reactivity effect in multisensory SJ or TOJ tasks or compared these effects between the two tasks. Since different neural mechanisms are believed to underlie TOJ and SJ tasks (Binder 2015, Basharat et al., 2018), a direct comparison could provide valuable insights into the mechanisms differentiating these processes.

Collecting confidence data in these tasks has several additional benefits. Prior research indicates that individuals are aware of their certainty in temporal judgments (Akdoğan & Balcı, 2017; Keane et al., 2015), allowing for the construction of psychometric functions based on confidence ratings (Allan, 1975). This approach facilitates an in-depth examination of how confidence influences TOJ and SJ task parameters. A previous study found a correlation between TBWs measured via TOJ and SJ tasks (Bedard & Barnett-Cowan, 2016), but it remains unclear whether this relationship extends to confidence-derived parameters. Confidence ratings have also been shown to reduce response variability, likely due to the reactivity effect (Yi & Merfeld, 2016), raising the possibility of training interventions that refine temporal perception. This is particularly relevant for older adults, who exhibit broader TBWs linked to increased fall risk (Mahoney et al., 2014; Setti et al., 2011).

In this study, we investigate whether the SJ and TOJ tasks exhibit reactivity, and explore the impact of confidence ratings on the TBW. Additionally, we examine correlations between confidence and perceptual parameters in these tasks. We hypothesize that both TOJ and SJ tasks will demonstrate a positive reactivity effect and that confidence-based psychometric functions will reflect previously observed response-based relationships (Bedard & Barnett-Cowan, 2016)). Lastly, we discuss the advantages and limitations of using Prolific for online data collection.

## GENERAL METHODS

### Participants

A total of 35 participants (15 males, 19 females, 1 preferred not to identify) aged 18 to 39 were included in the study. Of these, 29 were recruited via Prolific (www.prolific.com), and 9 were recruited as volunteers in person. Prolific consists of a database of 40,000 active participants and is designed for behavioral research scientists to conduct experiments online.

The online experiment was hosted on Pavlovia (www.pavlovia.org) which allows the experiment to run in the Firefox web browser. All participants reported having no auditory or visual disorders and no neurological conditions (Alzheimer’s, Stroke, Epilepsy, Parkinson’s disease, etc.) and gave their informed written consent in accordance with the guidelines of the University of Waterloo Research Ethics Committee. Participants on Prolific were remunerated $10 CAD for one hour of testing.

### Stimuli

Participants engaged in the study from their personal environment, using their own computer equipment. The experiment was accessible through Prolific for online recruitment and Pavlovia, with the stipulation that it be conducted via the Firefox web browser (version 107 or higher exclusively. Restrictions were placed to disallow the use of tablets and browsers other than Firefox to ensure uniformity in the experimental setup. Participants were instructed to use their computer’s speakers instead of headphones to simulate a unified auditory source. They were asked to maximize both their computer volume and screen brightness. Further, to optimize conditions for sensory processing, participants were advised to dim ambient lighting, minimize environmental noise, and maintain a consistent distance from their display, ideally an arm’s length away. Responses were recorded using the participants’ keyboards.

### Procedure

The study consisted of four randomized perceptual tasks. (1) SJ task without confidence ratings, (2) SJ task with confidence ratings, (3) TOJ task without confidence ratings and (4) TOJ task with confidence ratings. Each task began with participants focusing on a central fixation cross (visual angle of 1.5 degrees, when at arm’s length on a 15” laptop), designed to resemble a combination of a bull’s eye and crosshair, against a black background. Prior to each primary task, participants underwent six practice trials to familiarize themselves with the procedure. Stimuli presentation followed a randomized delay ranging between 1000 and 3000 milliseconds after the appearance of the fixation cross.

### Perceptual Tasks

In the auditory component of both the TOJ and SJ tasks, participants were exposed to a brief, auditory beep that was programmed to appear for 16 milliseconds with a frequency of 3500Hz. The visual stimulus consisted of a white circle, displayed 8 degrees below the fixation cross, programmed to appear for an equivalent duration of 16 milliseconds, against the same black background.The stimuli were presented with various Stimulus Onset Asynchronies (SOA), including 0, ±20, ±40, ±80, ±160, ±320, and ±640 milliseconds, allowing for a comprehensive assessment of temporal perception. A wide spread of SOAs, including the SOAs of ±320 and ±640 were used in order to include participants who may have wide TBWs, and to allow for comparison with previous and future studies in older adults, where higher SOAs are necessitated due to larger TBWs (Basharat et al., 2019). Because the experiment was not performed on a real-time operating system (RTOS), we cannot verify whether the stimuli appeared at the time programmed on the individual computers of participants, though we did validate stimulus presentation lengths and SOAS on home computers (Mac Mini with M1 chip). In the TOJ task, participants were tasked with identifying which stimulus, auditory or visual, appeared first by pressing 1 for visual and 2 for audio first responses on their keyboards. In the SJ tasks, participants had to identify whether the visual and auditory stimulus were simultaneous or not by pressing 1 for simultaneous and 2 for non-simultaneous on their keyboards. Emphasis was placed on accuracy over speed in the response. For the version of the tasks that included confidence ratings, participants quantified their certainty regarding their response on a scale from 0 to 100 percent after each decision by inputting a number on their keyboard. This scale was designed to capture the participants’ confidence, with the instruction to reserve extreme ratings for instances of absolute certainty or uncertainty. After feedback provided by in person volunteers, two participants’ confidence data was excluded due to errors in noting their confidence responses. The duration of each task was designed to be approximately 10 to 15 minutes, with an intermission that allowed participants to take a short break after each task. This break, lasting between 5 to 10 minutes, aimed to reduce the risk of fatigue impacting performance. The experimental design is summarized in Figure 1.

**Figure 1:**
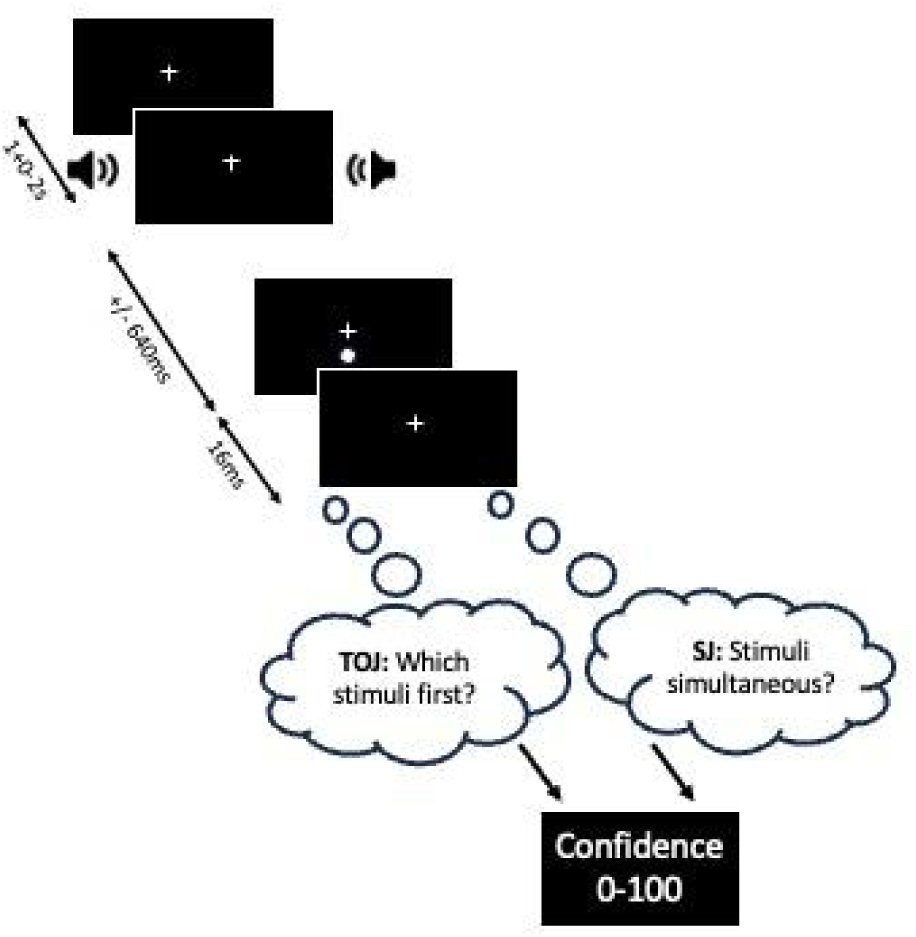
Schema of the TOJ and SJ tasks. Participants were presented with an audiovisual pair of stimuli which appear 1-3s following the appearance of a fixation cross. The visual stimulus can occur 0, ±20ms, ±40ms, ±80ms, ±160ms, ±320ms, ±640ms before (-) or after (+) the auditory stimulus. This figure depicts the auditory stimulus appearing before the visual stimulus. Both the auditory stimulus and the visual stimulus are 16ms long (adapted from Bedard & Barnett-Cowan 2016). Participants decide their answer for each task respectively (depicted in thought bubbles), and then are asked to rate their confidence in their decision.

### Data Analysis

Response data from participants was saved in .xlsx or .csv format. Raw recorded data was analyzed using Python 3.9.12 and using the python libraries scipy, numpy, pandas, sklearn and math. First, confidences were converted from “confidence in response” to “confidence that light came first” by subtracting 100 from the confidence if sound came first. This was to ensure that the confidence data would have a similar shape as the response data when fit to a psychometric function.

Psychometric functions were fitted to each participant’s response and confidence data as a function of SOA using Python 3.9.12, either using a sigmoidal logistic function for the TOJ task with additional lapse parameters to account for attentional lapses (Wichmann & Hill, 2001) bound maximally to 5% (Eq. 1) or a three-parameter Gaussian function also with additional lapse parameters bound maximally to 5% (Eq. 2).

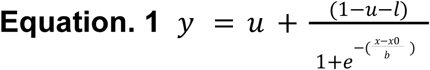

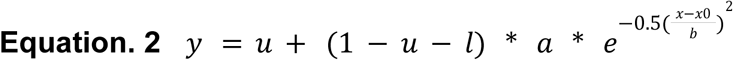

For both curves, *x0* represents the point of subjective simultaneity (PSS), which is the point at which the participants perceive the two stimuli as appearing simultaneous. The *b* parameter is the slope of the curve, which represents the sensitivity of the participants, or the minimum amount of time needed to discriminate temporal order or simultaneity. The parameters *u* and *l* represent the lapse parameters, and the parameter *a* represents the amplitude for Eq. 2. Since we are interested in the relative difference in precision between the two tasks, *b* is taken as a proxy for the TBW to avoid the differences in the literature concerning how to define the absolute size of the TBW. Initial values for fitting were set at *b* = 50 and *x0* = 0. Maximum iterations were set at 10,000 repeats. Fitting was performed via scipy’s curve fitting functions, which uses a non-linear least squares approach. After calculating best fit parameters, those participants whose data was not well estimated by the fit (r^2^ < 0.2) were excluded from further analysis.Metacognitive reactivity was measured as the difference in the b parameter (TBW) between the task with confidence ratings, and the task without confidence ratings, and was denoted as ΔTBW. Lastly, the confidence factor was obtained by dividing the b from the psychometric curve made from response data by the b from the psychometric curve made from confidence data, in line with Yi and Merfeld’s (2016) definition of the confidence factor. This parameter signifies the confidence of each individual participant. An underconfident participant will have a k>1 whereas an overconfident participant will have a k<1.

To test for normality of data, Shapiro-Wilk tests were used, and since all data collected was found to not be normally distributed, Wilcoxon signed-rank tests were used when testing differences between the TOJ and SJ tasks. To determine the effect and interaction of confidence and task type on the PSS and TBW (b-value), repeated-measures analysis of variance (ANOVA) with a 2 (task type) × 2 (confidence) design was used. Cohen’s d was used to measure effect size for Wilcoxon signed-rank tests, and Cohen’s f was used to measure effect size for ANOVAs. Due to non-normality of the data, Spearman correlations were used to assess the relationship between the TOJ and SJ tasks, and between tasks with and without confidence ratings.

## RESULTS

Participants completed both SJ and TOJ tasks with and without confidence ratings. Participants’ binary response data for the tasks with and without confidence ratings, as well as participant’s confidence data for the task with confidence ratings was fit to a Gaussian curve for the SJ task (Eq. 1), and a sigmoidal logistic curve (Eq. 2) for the TOJ task, equaling 6 curves in total. Average psychometric curves to response data and confidence data are shown in Fig 2. As noted above, both the Gaussian and sigmoidal logistic curves included a lapse parameter (Wichmann & Hill) to account for attentional lapses. From these curves, the PSS, and the TBW were determined. To get a measure of the confidence of each participant, the ratio of the b values from the confidence and response curves, named k, was taken. The descriptive statistics are summarized in Table 1 for the TBW (b-values), and in Table 2 for the PSS.

**Figure 2.**
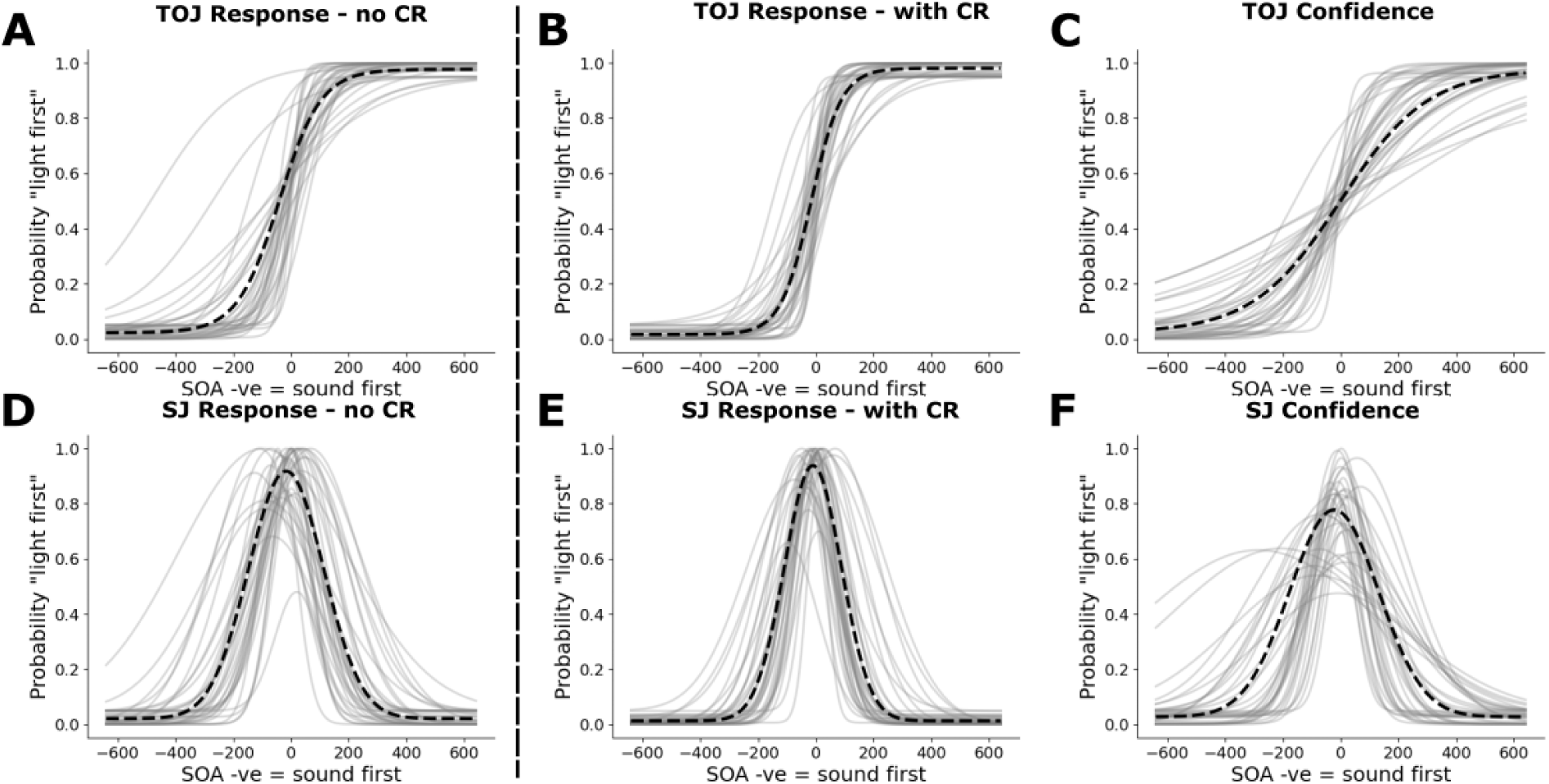
TOJ task (A-C): Sigmoidal logistic psychometric fits to average (thick line) and individual (thin line) to **A.** the TOJ task without confidence ratings to response data (PSS = -41.16, b/TBW = 72.24), **B.** the TOJ task with confidence ratings to response data (PSS = 17.12, b/TBW = 52.36) and **C.** the TOJ task with confidence ratings to confidence data (PSS = -23.08, b = 154.14). SJ task (D-F): Three-parameter Gaussian fits to average (thick line) and individual (thin line) to **D.** the SJ task without confidence ratings to response data (PSS = -17.12, b/TBW = 129.94), **E.** the SJ task with confidence ratings to response data (PSS = -10.52, b/TBW = 98.39) and **F.** the SJ task with confidence ratings to confidence data (PSS = -1.38, b = 151.53).

**Figure 3.**
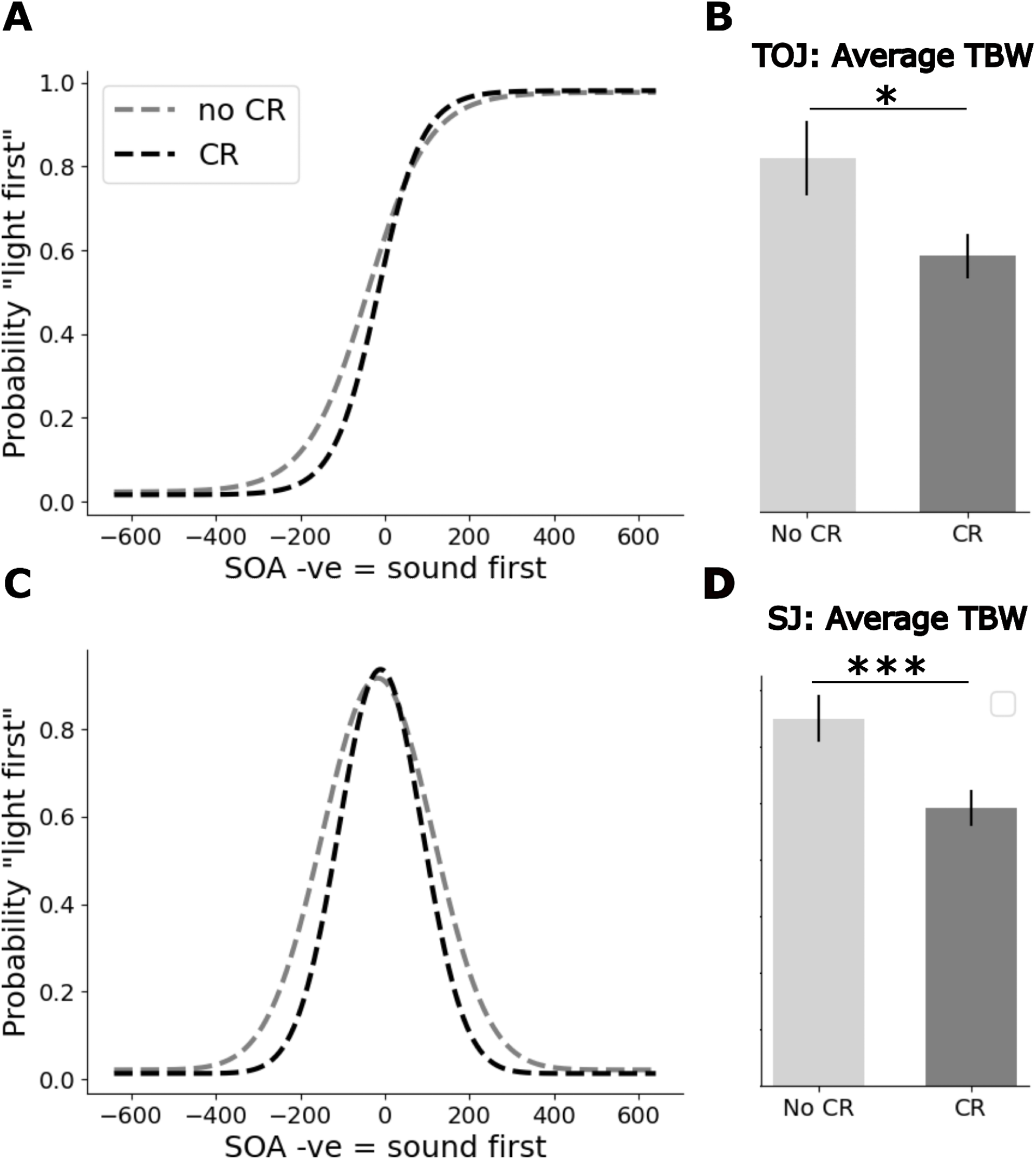
A. Average sigmoidal logistic fit to temporal order judgement data with (cr, black dashed line) and without confidence ratings (no cr, grey dashed line). **B.** Significant differences were found for the temporal binding window between the TOJ task with and without confidence ratings. **C.** Three parameter Gaussian fit to simultaneity judgement data with (cr, black dashed line) and without confidence ratings (no cr, grey dashed line). **D.** Significant differences were found for the temporal binding window between the SJ task with and without confidence ratings. Error bars are ±1 SEM

**Table 1.** Descriptive statistics for the TBW (b-value) for the response data and the b-value for the confidence data for all four tasks. CR = confidence ratings, Conf = Confidence

| Task | Mean | Median | Standard Dev. | Standard Err. |
| --- | --- | --- | --- | --- |
| TOJ without CR | 72.24 | 58.8 | 44.53 | 7.64 |
| SJ without CR | 129.94 | 116.2 | 48.44 | 8.31 |
| TOJ with CR | 52.36 | 53.17 | 26.22 | 4.5 |
| SJ with CR | 98.39 | 90.05 | 37.24 | 6.39 |
| TOJ CR Conf | 154.14 | 128.21 | 86.56 | 14.84 |
| SJ CR Conf | 151.53 | 117.31 | 102.93 | 17.65 |

**Table 2.** Descriptive statistics for the PSS for the response data and the b-value for the confidence data for all four tasks. CR = confidence ratings, Conf = Confidence.

| Task | Mean | Median | Standard Dev. | Standard Err. |
| --- | --- | --- | --- | --- |
| TOJ without CR | -41.16 | -23.6 | 93.34 | 16.01 |
| SJ without CR | -16.39 | -12.42 | 39.38 | 6.75 |
| TOJ with CR | -17.12 | -2.35 | 49.59 | 8.51 |
| SJ with CR | -10.52 | -3.57 | 33.45 | 5.74 |
| TOJ CR Conf | -23.08 | -4.41 | 66.41 | 11.39 |
| SJ CR Conf | -1.38 | -3.37 | 50.37 | 8.64 |

### Results from the TBW/b-value

The TBW was assessed with a 2 (SJ or TOJ task) x 2 (confidence vs. no confidence) repeated-measures ANOVA to determine the effects of task type, confidence, and their interactions. A significant within-subjects effect of task was found for the TBW (F = 69.4457, p < 0.0001, Cohen’s f = 1.41), where the SJ task showed a wider TBW than the TOJ task. Furthermore, a significant within-subjects effect of confidence was found for the TBW (F = 32.0650, p < 0.0001), where the task with confidence showed a narrower TBW than the task without confidence. The significantly narrower TBW for the tasks with confidence means that both the TOJ task and the SJ task exhibit positive reactivity. The interaction between task and confidence on the TBW was not significant (F = 1.8991, p = 0.1772), meaning that the impact of confidence does not significantly differ between the TOJ and SJ task. To see which of the two tasks were more strongly affected by the inclusion of confidence ratings, Cohen’s d were calculated for the difference between the TBW with and without confidence rating. These revealed an effect size of 0.54 for the TOJ task, and an effect size of 0.73 for the SJ task. Lastly, a Wilcoxon signed-rank test showed no significant differences between the b values of the confidence curve for the TOJ task and the SJ task (Z = 261, p=0.385).

A correlation between the TBW for the TOJ task without confidence ratings, and the change in TBW between TOJ tasks was found (r=0.62, *p*<0.001), where larger TBWs for the TOJ task without confidence ratings correlated with a larger change in TBW between TOJ tasks (see Fig 4A). This correlation was also found for the SJ task (r=0.55, *p*<0.001) where larger TBWs in the SJ task without confidence ratings correlated with a larger change in TBW between SJ tasks (see Fig 4B). K-means clustering of the change in TBW between tasks (reactivity) for the TOJ task versus the change in the TBW between tasks (reactivity) for the SJ task (Fig 4C) allowed for distinct “reactivity” groups to emerge. Distortion scores (inset on Fig 4C) confirmed an elbow at 4 clusters. These 4 clusters appear to visually represent participants who 1. Show more reactivity in the SJ task than the TOJ task (blue squares), 2. Show more reactivity in the TOJ task than the SJ task (green triangles), 3. Show low or even negative reactivity for both tasks (yellow circles) or 4. Show strong reactivity in the TOJ task and moderate reactivity in the SJ task (red diamonds).

**Figure 4.**
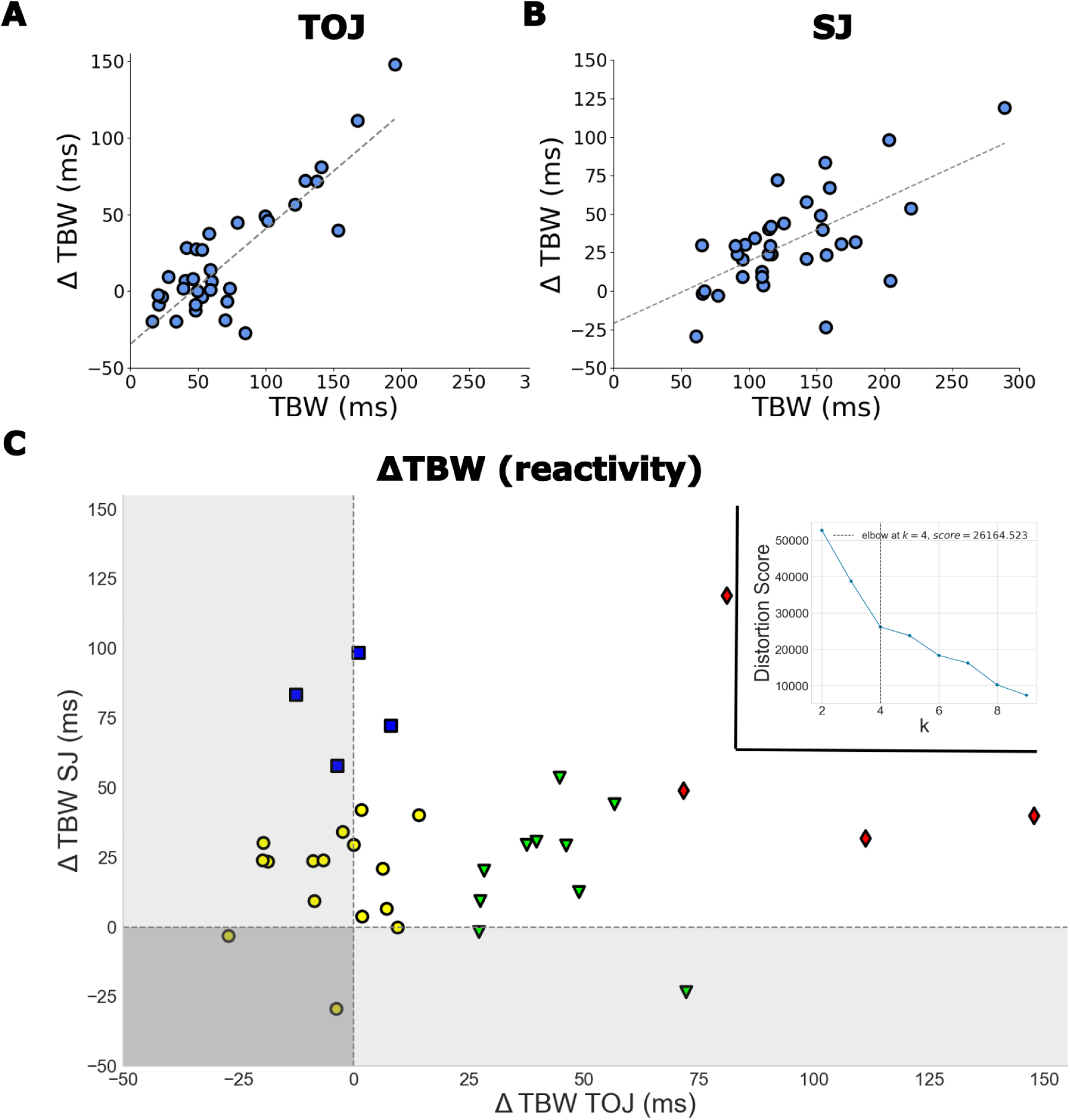
Linear Regressions (solid black line) of the TBW for the task without confidence ratings versus the difference in TBW between tasks for **A.** the TOJ task and **B.** the SJ task. Each blue dot represents one participant. **C.** Change in TBW between TOJ tasks versus change in TBW between SJ tasks. Values over 0 denote positive reactivity. Inset: K-means clustering distortion score reveals 4 clusters in the data, which are represented in C as the four different symbols and colors.

In line with previous research (Bedard & Barnett-Cowan, 2016), significant positive correlations were found for the TBW between the TOJ and SJ tasks, both in the tasks without confidence ratings (r=0.50, p<0.01) and with confidence ratings (r=0.42, p<0.05). Novel to this study, a significant positive correlation was also found between the b value of the confidence curve for the TOJ task and SJ task (r=0.72, p<0.001).

A Wilcoxon signed-rank test revealed a significant difference between the confidence factor k in the TOJ task and SJ task (Z=41, *p*<0.001, Cohen’s d =1.11). However, no correlation was found between more confident participants and narrower TBWs for the TOJ task both with (r=-0.127, p=0.467) and without (r=-0.119 p=0.494) confidence ratings. Nor was there a correlation between confidence and change in TBW between tasks (r=-0.120, p=0.490). Likewise, no correlation was found between more confident participants and narrower TBWs for the SJ task, both for the task with (r=-0.057, p=0.743) and without confidence ratings (r=0.076, p=0.664). However, a significant correlation was found between confidence and change in TBW between tasks for the SJ tasks (correlation=0.336, p<0.05).

As indicated in the Methods section, a confidence of >1 indicates underconfidence, whereas a <1 indicates overconfidence. Confidence for the SJ task was on average 1.59 (Median=1.23, S.D.=0.81, S.E.=0.14) and 2.95 for the TOJ task (Median=2.53, S.D.=1.43, S.E.=0.24). As shown in Fig 5, Wilcoxon signed-rank tests on the tasks with confidence ratings revealed that participants were significantly underconfident for the SJ task (Z=15, *p*<0.001, cohen’s d=1.04) as well as the TOJ task (Z=2, *p*<0.001, cohen’s d=1.93).

**Figure 5.**
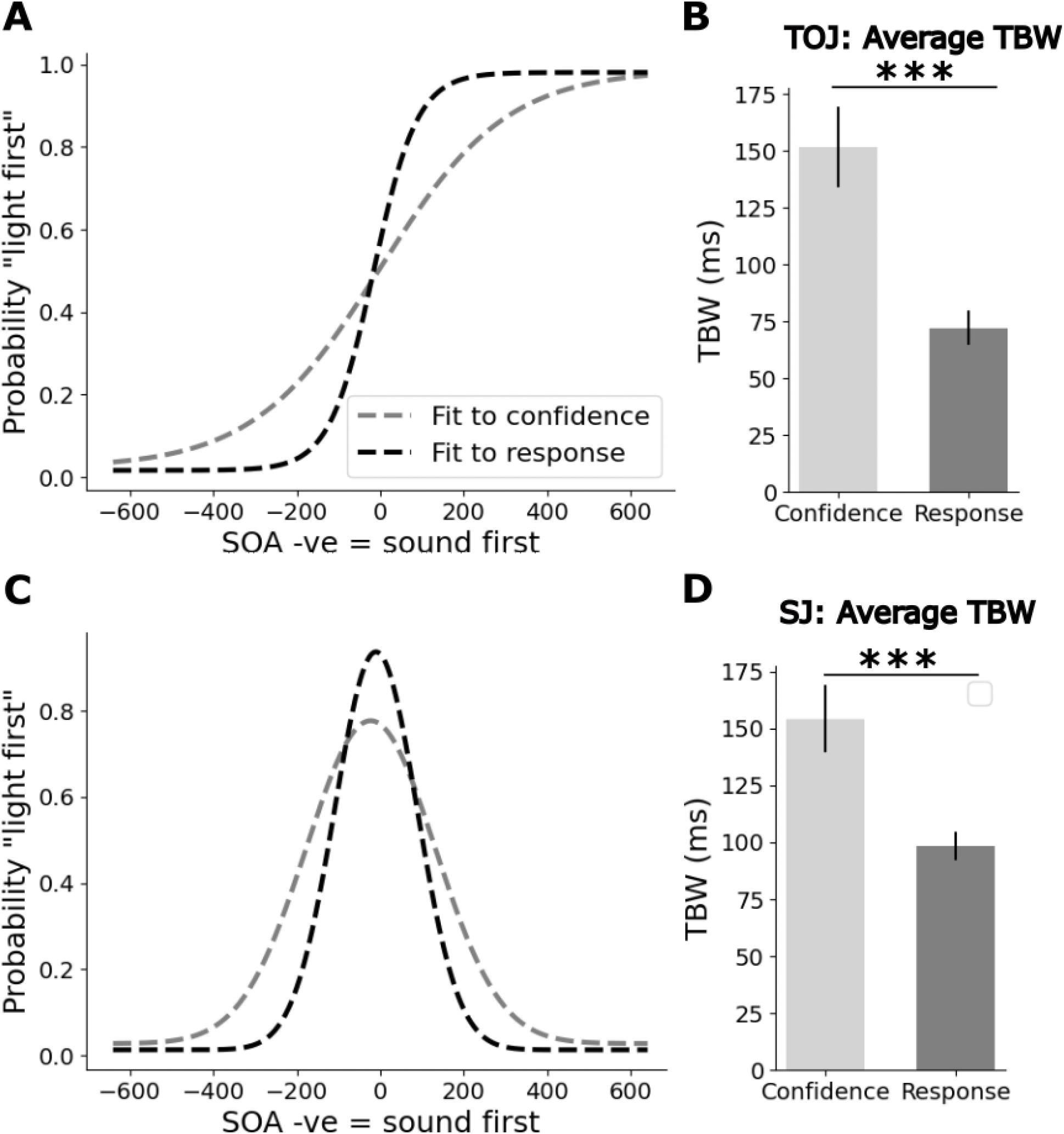
A. Average sigmoidal logistic fit to temporal order judgment data with confidence ratings to response data (fit to response, black dashed line) and to confidence data (fit to confidence, gray dashed line) showing significant underconfidence. **B.** Significant differences were found for the average b-value between the fit to confidence data and response data. **C.** Three parameter Gaussian fit to simultaneity judgment data with confidence ratings to response data (fit to response, black dashed line) and to confidence data (fit to confidence, gray dashed line) showing significant underconfidence. **D.** Significant differences were found for the average b-value between the fit to confidence data and response data. Error bars are ±1 SEM. **Results from the PSS**

Given that the experiment was not performed on an RTOS, true PSS values cannot be ascertained, instead relative differences between tasks were obtained. Differences in the PSS were assessed with a 2 (SJ or TOJ task) x 2 (confidence vs. no confidence) repeated-measures ANOVA to again determine the effects of task type, confidence, and their interactions.Here, no significant effects were found fo the task type (F=2.0795, p=0.158). A significant within-subjects effect of confidence was found (F=5.4162, p<0.05). No interaction was found between task and confidence for the PSS values. Lastly, Wilcoxon signed-rank test found no significant differences between the PSS from the confidence curves of the TOJ task (Mean=-23.08, Median=-4.41, S.D.=66.41, S.E=11.39) and the SJ task (Mean=-1.38, Median=-3.37,, S.D.=50.37, S.E=8.64) (Z=252, p=0.310).

Significant negative correlations were found between the PSS and confidence for the SJ tasks, both with (r= -0.5), p<0.001) and without confidence ratings (r=-0.37, p<0.05) with less confident participants resulting in a more visual leading PSS. No significant correlations were found for the TOJ task, both with (r=0.05, p=0.772) and without (r=-0.06, p=0.729) confidence ratings.

## DISCUSSION

In this study, our main objectives were to determine the strength and direction of the reactivity effect in the TOJ and SJ tasks with confidence ratings and to identify the factors influencing it. We found that both tasks exhibited a reactivity effect, demonstrating that positive reactivity can be observed in multisensory time perception tasks. The strength of this effect was influenced by the size of participants’ TBW in the “no confidence” condition, the specific task being performed, and, in the case of the SJ task, participants’ confidence levels. To our knowledge, this is the first study to examine the reactivity effect of confidence ratings in time perception tasks. The closest comparable study involved a time reproduction task in which participants self-corrected biases when incentivized (Brocas et al., 2018), though it did not explicitly investigate reactivity. Additionally, while we observed differences in reactivity between the SJ and TOJ tasks, we found a significant correlation between the perceptual estimates of confidence curve widths for both tasks. Furthermore, there was considerable heterogeneity in the strength and direction of reactivity across participants, who appeared to cluster into distinct groups. Lastly, we explored the relationships between confidence data from the SJ and TOJ tasks and discussed our experience using Prolific as a platform for participant recruitment.

For both the SJ and TOJ tasks, we found a strong correlation between TBW size in the “no confidence” condition and the strength of the reactivity effect. Participants with larger TBWs exhibited greater positive reactivity, with this effect being more pronounced in the TOJ task. A potential ceiling effect must be considered—if a participant’s TBW is already narrow before confidence ratings are introduced, there is limited room for improvement. This suggests that TBW size is a key factor in determining reactivity strength.

Our findings indicate that participants were underconfident in both tasks, a common trend in perceptual studies that may stem from attributing the same confidence rating to stimuli of varying difficulty (Stankov et al., 2012). We observed that participants were less confident in their responses for the TOJ task and exhibited lower reactivity compared to the SJ task. The difference in reactivity between tasks approached significance (p = 0.09), with effect sizes of 0.73 for the SJ task and 0.54 for the TOJ task. Participants tend to be underconfident in easy tasks and overconfident in more difficult ones, a phenomenon known as the “hard-easy effect” (Lamotte & Droit-Volet, 2017; Petrusic & Baranski, 2003). At first glance, this might suggest that the TOJ task is easier than the SJ task. However, prior research does not support this assumption. The TOJ task is considered a second-order judgment requiring an initial simultaneity decision before determining temporal order (Allan, 1975; Binder, 2015). Additionally, aging research suggests that temporal order discrimination declines while simultaneity discrimination remains stable (Bedard & Barnett-Cowan, 2016), implying that TOJ mechanisms depend on those underlying SJ. Furthermore, differences in decision-making processes (García-Pérez & Alcalá-Quintana, 2015) and increased sensory noise in TOJ tasks (Yarrow et al., 2014) suggest higher difficulty for TOJ. An fMRI study also indicated that TOJ requires greater cognitive resources than SJ (Binder, 2015). To our knowledge, the hard-easy effect was not investigated in TOJ or SJ tasks. Taken together, our results suggest that the hard-easy effect is not found in the TOJ or SJ task. Performance in the Raven’s progressive Matrix indicates that the difficulty of an item appears to affect reactivity - performance on easier items is higher when confidence ratings are included (Birney et al., 2017). Though we are dealing here with a more simple perceptual task, if we take for granted that the TOJ task is more difficult, perhaps then, the more difficult the task, the less reactivity we find.

### Neural Mechanisms underlying SJ and TOJ Tasks

An alternative explanation, independent of task difficulty, is that SJ and TOJ tasks engage different neural networks. The TOJ task recruits a more extensive network of brain regions (Binder, 2015), which may be differentially affected by metacognitive assessments. Supporting this, we found that confidence influenced reactivity in the SJ task—more confident participants showed greater reactivity—but this effect was absent in TOJ. Furthermore, while several participants exhibited positive reactivity in the TOJ task but not the SJ task, only two participants showed positive reactivity in the SJ task but not the TOJ task. Confidence also affected PSS in the SJ task but not TOJ, with more confident participants exhibiting more “visual leading” responses. These results not only align with prior findings that different neural mechanisms underlie SJ and TOJ tasks (Basharat et al., 2018; Bedard & Barnett-Cowan, 2016; Binder, 2015; Linares & Holcombe, 2014), but also appear to suggest that metacognitive assessments impact SJ and TOJ in different ways. In line with (Basharat et al., 2018) and (Love et al., 2013), we also found a significant difference in the TBW between SJ and TOJ tasks.

Moreover, although TBW values for SJ and TOJ tasks were correlated, reactivity in one task did not predict reactivity in the other. Our clustering analysis revealed four participant groups: those with greater reactivity in SJ but not TOJ (n=10), those with greater reactivity in TOJ but not SJ (n=5), those with minimal or no reactivity in either (n=16), and those with strong reactivity in both (n=4). These findings suggest substantial individual differences in reactivity. Though Binder (2015) did not look at whether heterogeneity of participants’ responses clustered into distinct groups, they did have to tailor their experiments using individual threshold of simultaneity for each participant, due to high variability among participants.

### Implications for Clinical Applications

The reactivity effect may have practical applications beyond metacognitive assessment. If metacognitive assessments can narrow the TBW, they could potentially aid clinical populations with inefficient multisensory processing. Older adults typically have larger TBWs, contributing to age-related issues such as increased fall risk (Mahoney et al., 2014; Setti et al., 2011). Since younger adults (like those in our study) generally have narrower TBWs, a ceiling effect may limit their reactivity. However, older adults may exhibit stronger reactivity effects due to greater room for TBW improvement. Prior studies have explored feedback-based TBW narrowing, though results have been mixed (O’Brien et al., 2020; Powers III et al., 2016; Setti et al., 2014), potentially due to participant fatigue. Future research should explore whether reactivity effects can be harnessed in training paradigms for clinical populations.

### Correlation of Confidence Parameters between SJ and TOJ

Consistent with Bedard & Barnett-Cowan (2016), we found that while the PSS between TOJ and SJ tasks was not correlated, the TBW for both tasks showed a significant correlation. Since phenomena such as blindsight, along with extensive neural and behavioral evidence, suggest that confidence is encoded in distinct brain regions separate from decision-making circuitry (see Grimaldi et al., (2015) for a review), it was unclear whether psychometric parameters derived from confidence data would also correlate. However, data from Keane et al., (2015) suggest that temporal judgments and confidence share common informational sources, as trial-by-trial variability distinguished high- and low-confidence trials. In line with this, we found a strong correlation in the b-value derived from confidence data in both the SJ and TOJ tasks. This suggests shared neural mechanisms underlying confidence across these tasks, despite differences in the reactivity effect and overall confidence levels. Although the TOJ task recruits additional left-hemisphere regions (Binder, 2015), both the TOJ and SJ tasks activate common areas, particularly within the bilateral fronto-parietal network, which is implicated in spatial selective attention. Notably, the correlation between TOJ and SJ task parameters appears to diminish with aging (Bedard & Barnett-Cowan, 2016). Future studies should further investigate this relationship in older adults, particularly in the context of confidence-related measures.

### Limitations

Several limitations should be noted. First, since we did not compare unisensory to multisensory tasks, we cannot determine how multisensory stimuli influenced reactivity. Second, the wording of the confidence prompt (e.g., explicitly asking about “confidence”) may have induced positive reactivity by making participants more aware of their self-perceived ability (Double & Birney, 2019a). Future studies should explore variations in task wording and assess self-confidence to better understand interindividual differences. Lastly, our study was conducted online using Prolific, and participants did not use a real-time operating system (RTOS). This prevents us from making absolute PSS judgments within each task, though we could reliably compare PSS estimates between tasks.

### Online Data Collection with Prolific

Our experience using Prolific for online perceptual tasks had both advantages and drawbacks. The platform enabled rapid recruitment, with access to over 22,000 potential participants after applying exclusion criteria. Compared to traditional university-based samples, Prolific provides a more diverse participant pool, including a large older adult population, making it useful for aging research. However, the lack of an RTOS introduces timing variability, potentially affecting SOA accuracy. Our PSS estimates suggest that SOAs were not presented veridically, as both SJ and TOJ tasks yielded negative PSS values. This does not impact our main goal of comparing TBW between tasks but warrants caution for researchers conducting time-sensitive perceptual studies online. Pilot analyses indicated that restricting participants to Mac and Linux users reduced PSS lag. Despite concerns about data quality in crowdsourced studies, prior research suggests that Prolific participants provide more reliable responses than those on other platforms (Douglas et al., 2023). Moreover, our TBW estimates were comparable to those obtained from in-person RTOS studies. For studies focusing on relative psychometric differences, Prolific offers a viable and efficient alternative to traditional lab-based research.

## Acknowledgments

Supported by a Natural Sciences and Engineering Research Council of Canada (NSERC) Discovery Grant (RGPIN-03977-2020) awarded to MB-C. CS was supported by funding from the Gender Equality Commission of the Faculty of Science with mentorship from Dr. Denzinger and Prof. Dr. Mallot. We also would like to acknowledge Matt Harvey for coding the experiment for Pavlovia.

